# The fallacy of identification by neutralization in the light of cytopathology nonproducing enterovirus strains

**DOI:** 10.1101/173419

**Authors:** Moses Olubusuyi Adewumi, Temitope Oluwasegun Cephas Faleye, Johnson Adekunle Adeniji

**Affiliations:** Department of Virology, College of Medicine, University of Ibadan, Ibadan, Oyo State, Nigeria.; Department of Microbiology, Faculty of Science, Ekiti State University, Ado Ekiti, Ekiti, State, Nigeria.; WHO National Polio Laboratory, University of Ibadan, Ibadan, Oyo State, Nigeria.

**Keywords:** Nigeria, Enterovirus, Recombination, CV-A24/PV2, Cytopathology

## Abstract

We describe the characterization of an enterovirus isolate recovered from untreated raw sewage in Ibadan, southwest Nigeria in 2010. The isolate was neutralized by specific antisera and consequently identified as Echovirus 7 (E7). Subsequent molecular characterization showed the isolate to be a mixture of E7 and Coxsackievirus A24 (CV-A24) thereby suggesting the CV-A24 strain to be non-cytopathology producing. Further molecular analysis suggested that the CV-A24 might have recombined with a Sabin poliovirus 2 (PV2) in its non-structural region. This is the first description of a Nigerian CV-A24 strain.

---

Enteroviruses (EVs) are members of the family *Picornaviridae*, order *Picornavirales*. There are 13 species in the genus and Enterovirus Species C (EV-C) contains the prototype member of the genus; Poliovirus. EVs have a 27 – 30nm diameter naked icosahedral capsid which encapsidates a 5^*l*^ protein-linked, positive-sense, single-stranded, ~7.5kb, RNA genome. The genome is flanked on both ends by 5^*l*^ and 3^*l*^ untranslated regions (UTRs) and has a single open reading frame (ORF) between the UTRs. The ORF is translated into a polyprotein that is self-cleaved into three smaller proteins (P1, P2 and P3) and subsequently into 11 protein products. VP1-VP4 (from P1) are the structural proteins which form the capsid. While 2A-3D (from P2 & P3) are nonstructural proteins crucial for virus replication.

Classically enterovirus detection is by isolation in cell (usually RD and L20B) culture (WHO, 2003, 2004), although this is fast being replaced by cell-culture independent strategies (WHO, 2015). Also, identification was classically by neutralization using polyclonal antisera. However with an association shown between VP1 sequence data and enterovirus serotypes VP1 amplification and sequencing is now being used for enterovirus identification (Oberste *et al*., 2001, 2003).

In June 2010 we isolated an enterovirus in RD cell culture. The isolate was from a sample of untreated raw sewage collected in Ibadan, southwest Nigeria. The sample was collected as part of an environmental surveillance study conducted by our group (unpublished). It was subsequently concentrated and enterovirus recovered from it by culture in RD cell line as previously detailed (Adeniji and Faleye, 2014). The RD cell culture isolate was subjected to neutralization (WHO, 2004; Adeniji and Faleye, 2014) using a panel of antisera provided by RIVM, Netherland and identified as Echovirus 7 (E7). On passage in L20B cell line, the isolate did not show cytopathology (CPE). Interestingly, on subjection to both the Panenterovirus and Panpoliovirus real-time reverse transcriptase polymerase chain reaction (rRT-PCR) assay in use by the Global Polio Laboratory Network and previously detailed in Adeniji and Faleye, (2014a), the isolate was positive for both but with variations. While the Panenterovirus assay was clearly positive with an early (≤ 30 cycles) Ct value, the Panpoliovirus assay was controversially positive with a late (≥ 32 cycles) Ct value. The result of the Panpoliovirus rRT-PCR assay did not improve despite repeated cycles of passage in RD cell line and subsequent Panpoliovirus rRT-PCR screen.

In, this study we attempt to better characterize this E7 isolate in a bid to determine the reason an RD cell culture positive but L20B negative isolate is showing late positivity with very high Ct value on subjection to the Panpoliovirus rRT-PCR assay. The algorithm followed in this study is depicted by Figure 1.

**Figure 1:**
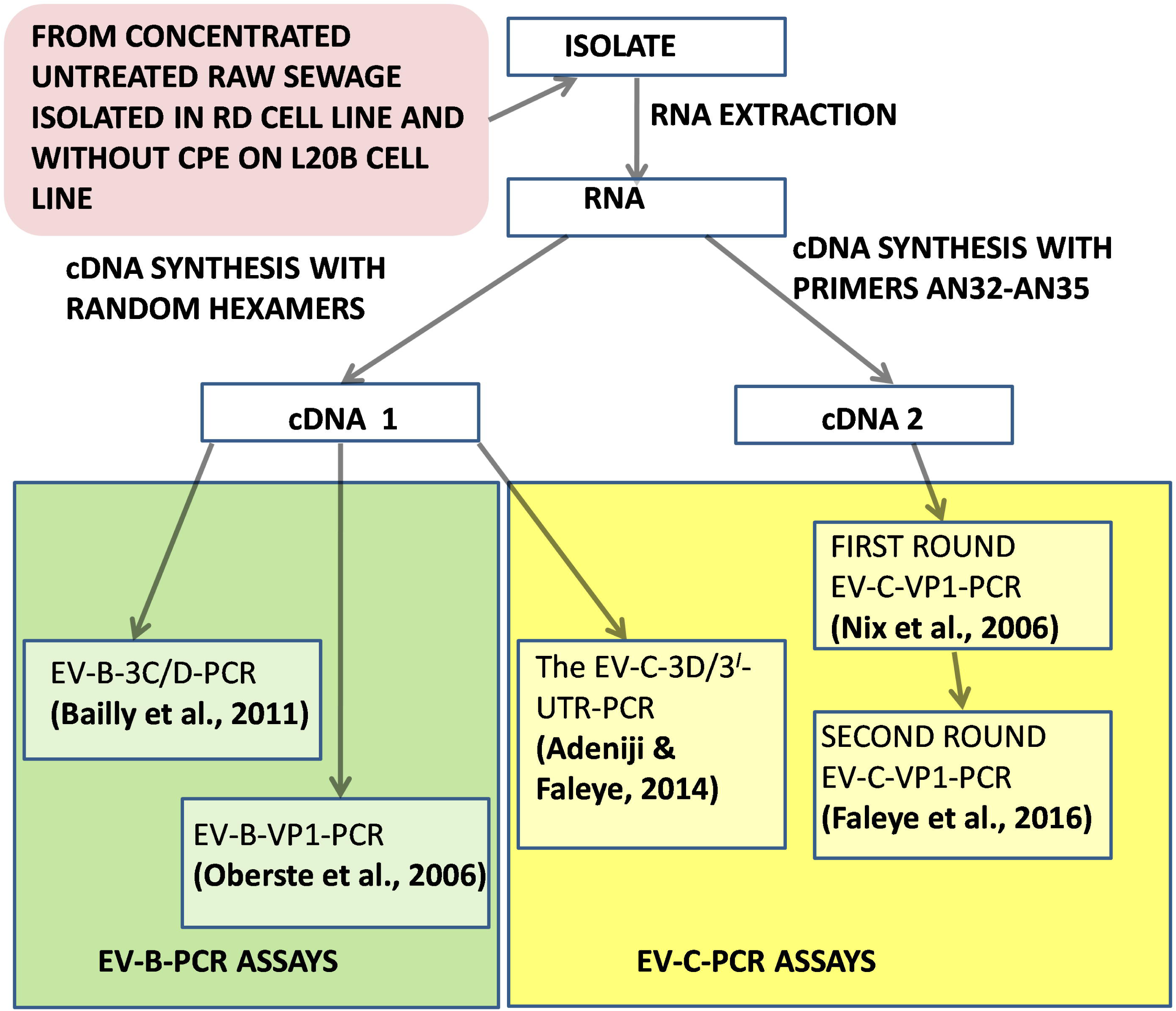
A schematic representation of the algorithm followed in this study.

RNA was extracted using the JenaBioscience RNA extraction kit (Jena Bioscience, Jena, Germany). Using the Script cDNA synthesis kit (Jena Bioscience, Jena, Germany) two different cDNAs were made (Figure 1). The basic difference was that the first cDNA (cDNA 1) was done using random hexamers while the second cDNA (cDNA 2), was done using primers AN32, AN33, AN34 and AN35 (Faleye et al., 2016). A Veriti thermal cycler (Applied Biosystems, California, USA) was used for thermal cycling.

Four (two each for EV-B & EV-C) different polymerase chain reaction (PCR) assays were done. For each species one of the PCR assays targeted the VP1 (structural region) and the other the nonstructural (3C/D for EV-B and 3D/3^*I*^-UTR for EV-C) region (Figure 1). All the four assays (EV-B-3C/D-PCR, EV-B-VP1-PCR, EV-C-3D/3^*I*^-UTR-PCR and EV-C-VP1-PCR) were two-step RT-PCR assays except for EV-C-VP1-PCR which is an RT-semi-nested PCR (RT-snPCR) assay (Faleye et al., 2016). All amplicons generated were sequenced and isolate identity determined using the enterovirus genotyping tool (Kroneman et al., 2011). Multiple sequence alignments and phylogenetic trees were done using the default settings of the CLUSTAL ***W*** program in MEGA 5 software (Tamura et al., 2011) as previously described (Faleye et al., 2016). The sequences obtained from this study have been deposited in GenBank with accession numbers KM264408, KM264416, MF535106 & MF535107.

The isolate was positive for all four assays. All four amplicons were successfully sequenced and exploited for virus identification. The, EV-B-3C/D-PCR result confirmed that the isolate contained a Species B enterovirus. The EV-B-VP1-PCR result confirmed that the EV-B in the isolate is E7, thereby confirming the results of the neutralization assay. The EV-C-3D/3^*I*^-UTR-PCR showed that the isolate also contained a Species C enterovirus and the EV-C-VP1-PCR showed that the EV-C contained in the isolate is Coxsackievirus (CV) A24 (CV-A24).

Considering the significance of EV-Cs to poliovirus eradication particularly with respect to the emergence of circulating Vaccine Derived Polioviruses (cVDPV), only the EV-Cs were subjected to phylogenetic analysis. Figure 2 shows that, with respect to the VP1 sequence, the CV-A24 strain detected in this study belongs to a lineage that has only been described in sub-Saharan Africa till date. A BLAST search of the EV-C-3D/3^*I*^-UTR sequence (data not shown) showed it to be most similar to that of non-recombinant VDPV2s suggesting the CV-A24 might have recombined with a Sabin PV2.

**Figure 2:**
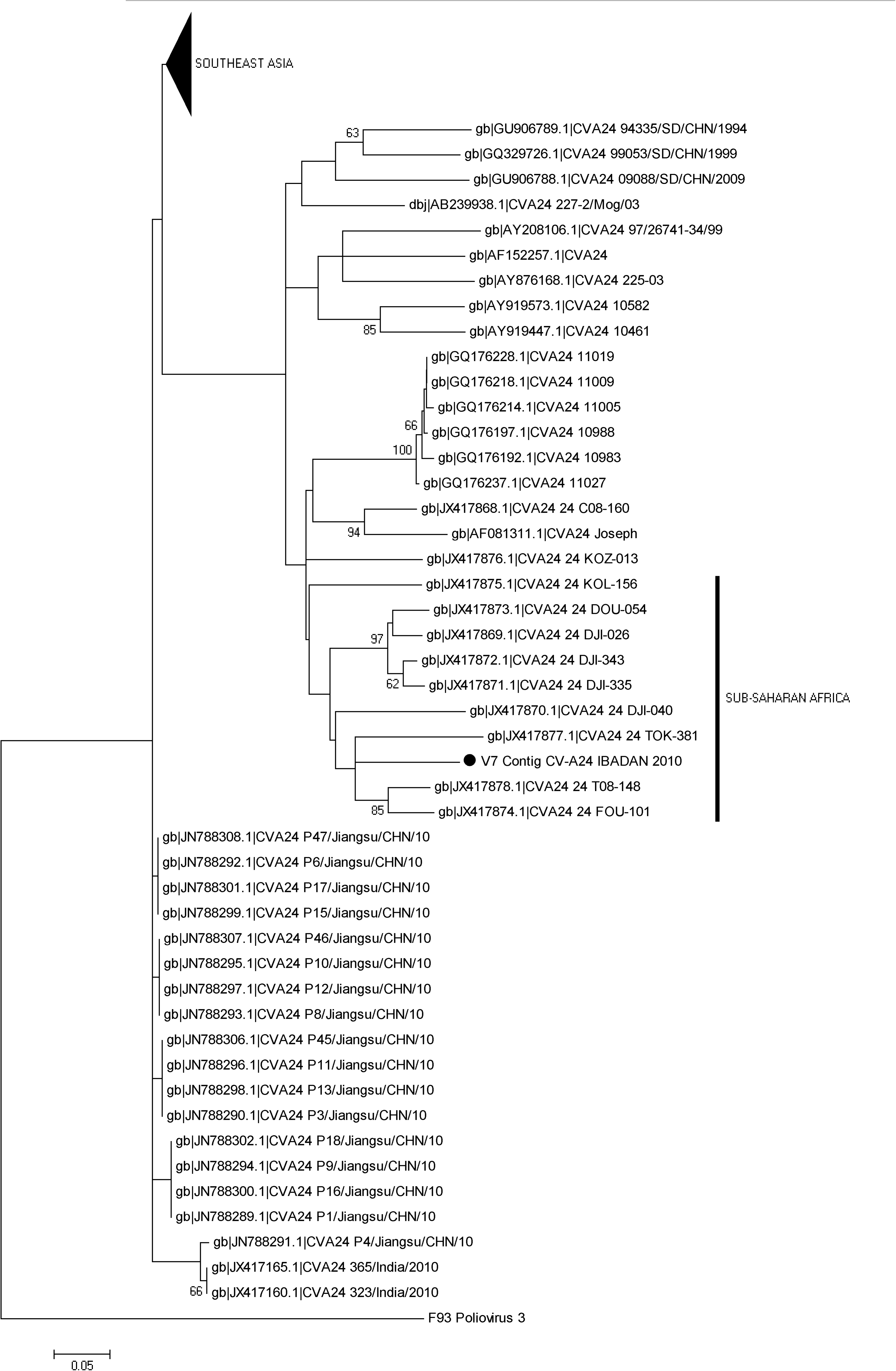
Phylogram of Coxsackievirus A24. The phylogram is based on an alignment of partial VP1 sequences. The newly sequenced strain is highlighted with Black circle. The GenBank accession number of the strains are indicated in the phylogram. Bootstrap values are indicated if > 50%. The labelled vertical bar is for ease of reference only.

In this study we attempted to determine the reason an already serotyped E7 that showed CPE on RD but not in L20B cell culture, is showing late positivity with very high Ct value (>32 cycles) on subjection to the Panpoliovirus rRT-PCR assay. We found that the presence of a CV-A24 that might have recombined with Sabin PV2 (CV-A24/PV2) in the isolate was the source of the signal. To the best of the authors’ knowledge, this is the first description of a Nigerian CV-A24 strain. The Panpoliovirus rRT-PCR assay was specifically designed to screen for the presence of poliovirus in isolates that had shown CPE in both RD and L20B cell lines in tandem (which is not the case with this isolate). The results of this study however show that the assay might also be valuable for detecting nonpolio enterovirus C (NPEV-C) members when used outside of its prescribed algorithm.

What is mostly found with recombinant EV-Cs is structural region of poliovirus origin and non-structural region of NPEV-C origin (PV/NPEV-C) and not the reciprocal (NPEV-C/PV). Why this is the case is currently not clear. It has however been shown (Jiang et al., 2007) that though such NPEV-C/PV recombinants might be replication competent, they tend to be less fit. Whatever the case, our finding of a possibly, CV-A24/PV2 recombinant, in this study, shows that such reciprocal recombinants might exist in nature, be circulating and even survive in the environment.

Though the isolate was a mixture of E7 and CV-A24, the fact that it was neutralized by antisera specific for E7 suggest that either the CV-A24 was very slow growing or was not replicating in RD cell line with evident CPE. In the light of Jiang et al., (2007), this is not totally strange. In fact, we have also recently isolated some EV-Cs that replicate in RD cell line without cytopathology (Adeniji et al., unpublished) and thus, further buttress the likelihood of such find. If indeed the CV-A24 does not produce CPE in RD cell line and was replicating alongside the CPE producing E7, it is crucial that the logic underlying enterovirus neutralization assays be revisited.

The concept of neutralization by specific antibodies is predicated on the assumption that any enterovirus present in the cell line of interest will produce CPE. The findings of this study question this assumption and shows it can sometimes be false. Although enterovirology is moving away from the classic technique of enterovirus identification by neutralization and depending more on molecular identification, a lot of studies are still dependent on the use of antibodies raised against enterovirus isolates. It is therefore essential to ensure that pure cultures intended for such purpose are truly pure and do not have any non-CPE producing strain(s) lurking unnoticed.

## ETHICS APPROVAL AND CONSENT TO PARTICIPATE

Enterovirus isolate was analysed in this study. The isolate was recovered from untreated raw sewage. Thus, this article does not contain any studies with human participants performed by any of the authors.

## ACKNOWLEDGEMENTS

We thank the WHO National Polio Laboratory in Ibadan, Nigeria for providing the cell lines used for isolating the enterovirus analysed in this study.

## AUTHOR CONTRIBUTIONS

1. Study Design (All Authors)
2. Sample Collection, Laboratory and Data analysis (All Authors)
3. Wrote, revised, read and approved the final draft of the Manuscript (All Authors)

## REFERENCES

Adeniji, J. A. and Faleye T. O. C. (2014a). Isolation and identification of enteroviruses from sewage and sewage contaminated water in Lagos, Nigeria. Food and Environmental Virology, 2014a, 6:75-86

Adeniji J.A., and Faleye, T.O.C., (2014b). Impact of Cell Lines Included in Enterovirus Isolation Protocol on Perception of Nonpolio Enterovirus Species C Diversity. J. Virol. Methods. 2014: 207: 238-247.

Bailly, J.-L., Mirand, A., Henquell, C., Archimbaud, C., Chambon, M., Regagnon, C., Charbonne, F., Peigue-Lafeuille, H., (2011). Repeated genomic transfers from echovirus 30 to echovirus 6 lineagesindicate co-divergence between co-circulating populations of the two human enterovirus serotypes. Infection, Genetics and Evolution 11,276–289.

Faleye, T.O.C., Adewumi, M.O., Kareem, S.A., Adesuyan, Y.O., Fapohunda, F.A., Fasanya, S.T., Jimeto, T., Lawrence, O.E., Obembe, A.A., Adeniji, J.A. (2016). The impact of a panenterovirus VP1 assay on our perception of the enterovirus diversity landscape of a sample. J Hum Virol. Retrovirol. 4(3):00134.

Jiang P, Faase JA, Toyoda H, Paul A, Wimmer E, Gorbalenya A.E. (2007) Evidence for emergence of diverse polioviruses from C-cluster coxsackie A viruses and implications for global poliovirus eradication. Proc Natl Acad Sci U S A 104: 9457–9462.

Kroneman A., Vennema H., Deforche K., v d Avoort H, Peñaranda S, Oberste MS, Vinjé J, Koopmans M. (2011). An automated genotyping tool for enteroviruses and noroviruses, Journal ofClinical Virology, vol. 51, no. 2, pp. 121–125.

Nix, W. A., Oberste, M. S., Pallansch, M. A. (2006) Sensitive, Seminested PCR Amplification of VP1 Sequences for Direct Identification of All Enterovirus Serotypes from Original Clinical Specimens. Journal of Clinical Microbiology, 44(8): 2698–2704

Oberste, M. S., Nix, W. A., Maher, K., Pallansch, M. A. (2003). Improved molecular identification of enteroviruses by RT–PCR and amplicon sequencing. Journal of Clinical Virology. 26(3), 375-377.

Oberste, M. S., Schnurr, D., Maher, K., al-Busaidy, S., Pallansch, M. A. (2001). Molecular identification of new picornaviruses and characterization of a proposed enterovirus 73 serotype. Journal of General Virology. 82(2), 409-416.

Oberste, M.S., Maher, K., Williams, A.J., Dybdahl-Sissoko, N., Brown, B.A., Gookin, M.S., Penaranda, S., Mishrik, N., Uddin, M. and Pallansch, M.A., (2006). Species-specific RT-PCR amplification of human enteroviruses: a tool for rapid species identification of uncharacterized enteroviruses. Journal of General Virology. 87, 119–128

Tamura K., Peterson D., Peterson N., Stecher G., Nei M. and Kumar S., (2011). MEGA5: molecular evolutionary genetics analysis using maximum likelihood, evolutionary distance, and maximum parsimony methods. MolBiolEvol 28:2731–2739.

World Health Organisation (2003) Guidelines for environmental surveillance of poliovirus circulation. Geneva.

World Health Organisation (2004) Polio laboratory Manual, 4^th^ edition, Geneva.

World Health Organisation (2015). Enterovirus surveillance guidelines: Guidelines for enterovirus surveillance in support of the Polio Eradication Initiative. Geneva.

